# PyrMol: A Knowledge-Structured Pyramid Graph Framework for Generalizable Molecular Property Prediction

**DOI:** 10.1101/2025.11.09.686426

**Authors:** Yajie Li, Qichang Zhao, Jianxin Wang

## Abstract

Expert pharmaceutical chemists interpret molecular structures through a sophisticated cognitive hierarchy, transitioning from local functional moieties to spatial pharmacophores and, ultimately, to macroscopic pharmacological and physicochemical profiles. However, conventional Graph Neural Networks frequently overlook this high-level chemical intuition by treating molecules as single-scale atomic topology. To bridge this gap between human expertise and computational inference, we propose PyrMol, a knowledge-structured pyramid representation learning framework. By constructing heterogeneous hierarchical graphs, PyrMol orchestrates information flow across atomic, subgraph, and molecular levels. Crucially, the subgraph level systematically integrates three complementary expert views comprising functional groups, pharmacophores, and retrosynthetic fragments. To harmonize these explicit domain priors with implicit computational semantics, we introduce an adaptive Multi-source Knowledge Enhancement and Fusion module that dynamically balances their complementarity and redundancy. A Hierarchical Contrastive Learning strategy further ensures cross-scale semantic consistency. Empirical evaluations across ten benchmark datasets demonstrate that PyrMol outperforms 11 state-of-the-art baselines. Furthermore, its “plug-and-play” versatility provides a framework-agnostic performance boost for existing GNN architectures. PyrMol thus establishes a principled data-knowledge dual-driven paradigm for AI-aided Drug Discovery, effectively leveraging domain knowledge to catalyze advances in molecular property prediction.

## I. Introduction

THE traditional drug discovery process is notoriously protracted and costly, often requiring over 15 years and exceeding $1 billion in investment for each successfully approved drug [1]. In response to these challenges, the integration of advanced artificial intelligence (AI) into drug discovery represents a significant paradigm shift, particularly in molecular property prediction as the cornerstone of pharmaceutical community [2]. AI-aided molecular property prediction facilitates the early stages of drug discovery, aiding in hit and lead compound screening, chemical structure optimization, and even novel molecular entity generation. This significantly shortens the R&D cycle, reduces costs, and minimizes human error [3]. These models capitalize on various molecular featurization methods, including sequence-based [4], structure-based [5], and image-based descriptors [6].

Notably, Graph Neural Networks (GNNs) stand out in learning molecular representations by propagating messages across chemical topologies [7]. Beyond foundational architectures (e.g., GCN [8], GIN [9], GAT [10]), specialized molecular networks like CMPNN [11] and AttentiveFP [12] have been explicitly tailored to model atom-bond interactions and extract higher-level substructural semantics, yielding superior predictive efficacy across standard evaluation datasets [13], [14], [15]. However, the performance of the aforementioned end-to-end models is often constrained by the limited availability of labeled molecular data. To address this bottleneck, pretraining frameworks (e.g., MolFCL [16], Himol [17], and KPGT [18]) have emerged as a pivotal school of thought to mitigate the reliance on extensive annotations, achieving remarkable progress by capturing rich structural semantics from unlabeled data.

Furthermore, expert chemists decode the causal relationships between molecular structures and properties through the identification of chemically significant substructures, rather than viewing molecules merely as low-level atom-bond networks [16]. Consequently, considerable research efforts [17], [18], [19], [20], [21] have pivoted towards incorporating chemical fragments to augment the predictive power of GNNs. Early substructure-enhanced methods typically rely on a single extraction paradigm. For instance, Hajiabolhassan et al. [20] proposed FunQG, a graph coarsening framework based specifically on functional groups to mitigate oversmoothing. Similarly, Jiang et al. [19] designed PharmHGT to extract vital substructures primarily derived from chemical reaction rules. While these methods have made strides in capturing localized semantics, their reliance on a single perspective inevitably creates an “information silo”, inherently limiting the model’s ability to capture the multi-view biochemical nature of molecules.

Fundamentally, distinct molecular properties are governed by different chemical dimensions. A comprehensive molecular profile requires integrating diverse expert knowledge: functional groups [21] for local physicochemical traits, pharmacophores [22] for spatial binding potentials, and retrosynthetic fragments [23] for macroscopic structural modularity. Crucially, their relative contributions vary significantly across downstream tasks; for instance, pharmacophores may dominate target affinity predictions, while functional groups contribute to analyse solubility. Therefore, predictions of different molecular properties are often reliable inferences derived from a combination of analyses from multiple perspectives.

To overcome this single-perspective limitation, recent approaches have explored incorporating diverse subgraphs. For example, Kengkanna et al. [24] investigated the applicability of multiple subgraph views, such as junction trees and functional groups, within the MMGX framework. However, despite aggregating multiple perspectives, MMGX predominantly rely on naive aggregation techniques, failing to account for the intricate complementarity and inherent redundancy among heterogeneous domain knowledge, and struggling to dynamically align diverse chemical semantics. Consequently, this type of models often exhibit suboptimal generalizability across complex molecular property prediction tasks.

To break through this bottleneck and bridge the gap between sophisticated human expertise and computational inference, we propose PyrMol, a knowledge-structured pyramid graph framework for generalizable molecular property prediction. The main contributions of our work are as follows:

- We propose PyrMol, an end-to-end heterogeneous pyramid framework that orchestrates message passing across atomic, subgraph, and molecular levels. Its core topological design, Pyramid Molecular Graph, serves as a “plug-and-play” module, explicitly leveraging domain knowledge to comprehensively capture multi-scale molecular representations.
- We design an adaptive multi-source knowledge enhancement and fusion module to seamlessly integrate disparate streams of chemical expertise. By dynamically balancing knowledge complementarity and redundancy, this module effectively mitigates the poor generalization and robustness issues stemming from single-perspective representations.
- We develop a hierarchical contrastive learning strategy across multiple representation scales (atoms, subgraphs, and molecules). This cross-scale alignment enforces semantic consistency and maximizes the discriminative power of the learned embeddings.

Extensive experiments demonstrate that the end-to-end PyrMol outperforms existing molecular graph neural networks and subgraph-enhanced methods, achieving results competitive with pretraining models. Additionally, our proposed pyramid molecular graph further improve the performance of GNNs and subgraph-enhanced methods. In summary, PyrMol provides a powerful and adaptable framework for molecular representation learning, contributing to more generalizable AI-driven drug discovery.

## II. Method

As depicted in Fig.1, the framework of PyrMol contains three core components. First, we construct a Heterogeneous Pyramid Molecular Graph (Fig.1.A) to hierarchically abstract a molecule into Atom-, Sub-, and Mol-level topologies. Second, a knowledge-guided heterogeneous Message Passing Neural Network (MPNN) with an adaptive fusion module (Fig.1.B) is deployed to update multi-scale representations. This module dynamically integrates multi-source knowledge into a unified Sub-Fuse embedding, aligning its dimension with Atom-and Mol-embeddings for cross-scale interactions. Third, a hierarchical contrastive learning strategy (Fig.1.C) mutually regularizes and aligns semantic representations across the three levels. Ultimately, the aligned features are concatenated for downstream property prediction via a fully connected network.

**Fig. 1.**
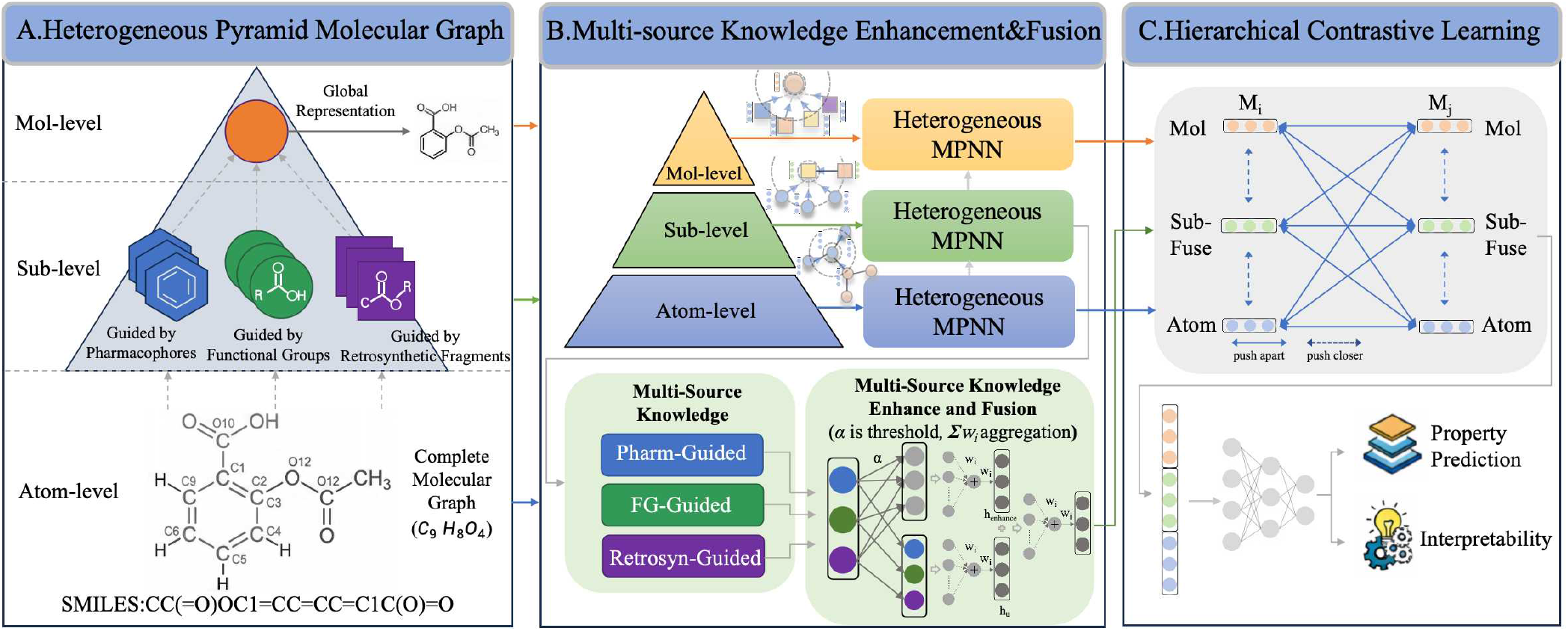
Illustration of Knowledge-Structured Pyramid Graph Framework(PyrMol). PyrMol contains three main modules: A. Heterogeneous Pyramid Graph; B. Multi-Source Knowledge Enhance and Fusion; C. Hierarchical Contrast.

### A. Heterogeneous Pyramid Message Passing Neural Network

#### 1) Pyramid Molecular Graph Construction

To explicitly guide GNNs in capturing hierarchical substructures, we construct a heterogeneous Molecular Pyramid Graph comprising three topological levels (Fig.1.a). The base level consists of atom nodes *V*_*A*_ and bond edges *E*_*A*_, representing the fundamental atomic topology. The middle level encompasses subgraph nodes 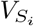 (representing key structural or functional motifs) and their relational edges. Specifically, 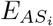 connects atoms to their corresponding subgraphs to direct explicit bottom-up feature aggregation, while 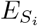 connects adjacent or overlapping subgraphs within the *i*-th knowledge type. The top level contains a single global molecular node *V*_*M*_ connected to all subgraphs via 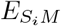, designed to aggregate localized semantics into a comprehensive global representation.

To populate the middle level, we integrate three complementary types of domain knowledge—functional groups (*F*), pharmacophores (*P*), and retrosynthetic fragments (*R*)—to parse molecules into distinct meaningful fragments. Specifically, we utilize RDKit to extract chemical functional groups (*V*_*F*_); employ the ErG algorithm [22] to identify biophysical pharmacophoric subgraphs (*V*_*P*_, encompassing six non-covalent interaction potentials such as H-bond donors/acceptors); and apply predefined BRICS rules [23] to generate retrosynthetically guided reaction fragments (*V*_*R*_). Consequently, the complete pyramid graph is formalized as *G* = {*V, E*}, where *V* = {*V*_*A*_, *V*_*P*_, *V*_*F*_, *V*_*R*_, *V*_*M*_} and *E* = {*E*_*A*_, *E*_*P*_, *E*_*F*_, *E*_*R*_, *E*_*AP*_, *E*_*AF*_, *E*_*AR*_, *E*_*P M*_, *E*_*F M*_, *E*_*RM*_}. Each node features are combined chemical features(detailed in Supplementary Table I) and computational features(initialed by Mol2Vec[25]). Initial node features are rapidly computed *in silico* via RDKit.

#### 2) Heterogeneous Message Passing Neural Networks

To endow the model with a robust baseline understanding of chemical semantics prior to hierarchical message passing, we initialize the intrinsic node attributes utilizing Mol2Vec embeddings [25]. It is imperative to note that within the PyrMol framework, these external embeddings serve strictly as static initial node features *F*_*i*_ rather than requiring a computationally expensive pretraining phase. Given the inconsistent dimensionality of node features across different types in heterogeneous graphs, PyrMol employs Full Connect Neural Networks(FCNNs) to standardize node features to *k* dimensions.

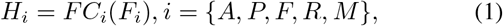

where 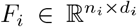 is the initial feature matrix for the *n*_*i*_ nodes of class *i* and *FC*(*) is FCNNs.

The architectural selection of our heterogeneous MPNN is explicitly driven by the varying nature of chemical message passing across scales. At the atomic and substructural levels, the heterogeneous graph formed by atom nodes *V*_*A*_ and subgraph nodes (*V*_*P*_, *V*_*R*_, *V*_*F*_) undergoes feature updates via GraphSAGE [26]. This leverages its inductive learning capability to efficiently aggregate localized topological environments and intra-subgraph connectivity. In contrast, at the molecular level, a Graph Attention Network (GAT) [10] is utilized to aggregate signals from these diverse substructure nodes to the global molecular node *V*_*M*_. This design choice is motivated by the chemical intuition that different substructures (e.g., a critical pharmacophore versus a generic carbon backbone) contribute unequally to the global molecular property, allowing the attention mechanism to dynamically learn these differential contributions.Formally, on each step *l* ∈ {0, 1, …, *L*} of the message passing phase, the hidden state 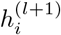 of the *i*-th node on the GraphSAGE layer is updated as follows:

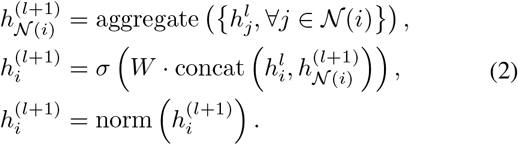

The hidden state of the molecular node 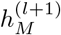 on the GAT layer is updated as follows:

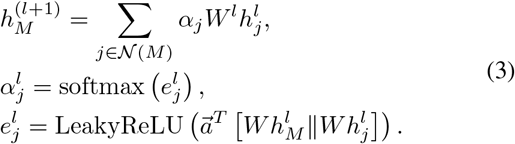

where *α*_*j*_ is the attention score between molecular node *M* and subgraph node *j*, and *a*^*T*^ is a learnable vector.

After *L* rounds of message passing, we have obtained the node features *H*^*L*^ = {*H*_*A*_, *H*_*P*_, *H*_*F*_, *H*_*R*_, *h*_*M*_}. By performing mean pooling on *H*_*i*_, *i* ∈ {*A, P, F, R*} respectively, the atom feature *h*_*A*_ ∈ ℝ^*k*^, functional group feature *h*_*F*_ ∈ ℝ^*k*^, pharmacophore feature *h*_*P*_ ∈ ℝ^*k*^, and retrosynthetic fragment feature *h*_*R*_ ∈ ℝ^*k*^ are obtained.

### B. Multi-source Knowledge Enhancement and Fusion Module

As shown in Fig.1.b., recognizing that disparate sources of expertise contribute unequally to molecular properties, we introduce a Multi-Source Knowledge Enhancement and Fusion Module. This module is designed to integrate divergent feature representations from various knowledge sources, highlighting the critical common features while orchestrating their integration in a principled manner.

Taking *h*_*F*_ ∈ ℝ^*k*^, *h*_*P*_ ∈ ℝ^*k*^, and *h*_*R*_ ∈ ℝ^*k*^, we calculate the similarity between pairs of them.

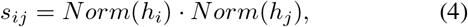

where *Norm*(*) means the normalization function, · is the dot product, and *i, j* ∈ {*F, P, R*}; *i* ≠ *j*.

Then we check if the similarity *s*_*ij*_ is greater than threshold *α* = 0.3. If it is, we generate their mutual key feature vector *k*_*ij*_ ∈ ℝ^*k*^ that are essential for them.

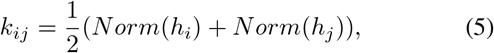

where the key feature matrix *K* ∈ ℝ^3×*d*^ is composed by {*k*_*F P*_, *k*_*F R*_, *k*_*P R*_}.

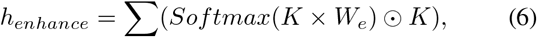

where *W*_*e*_ ∈ ℝ^*k*×1^ is a trainable weight and ⊙ is element-wise dot product (hadamard product). Then, we obtain the union feature vector of knowledge according to Eq.7

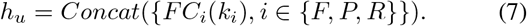

Finally, through learnable weight *W*_*f*_ ∈ ℝ^2*k*×*k*^ to balance flexibly union and enhance the characteristic, a fuse-level embedding is generated as follows.

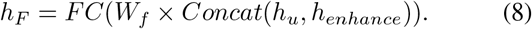

By mutually regularizing three streams of explicit domain priors (*V*_*F*_, *V*_*P*_, *V*_*R*_) with the implicit continuous computational semantics, this module establishes a powerful data-knowledge dual-driven paradigm. This principled synergy between discrete symbolic rules and data-driven representations dynamically highlights consensus features and effectively dismantles information silos, yielding a robust and comprehensive molecular embedding.

### C. Hierarchical Contrastive Learning Strategy and Prediction

Motivated by the goal of amplifying the model’s discriminative ability and unlock its strong expressive potential, we develop a hierarchical contrastive learning strategy. Concretely, the PyrMol model, as mentioned, yields three-floor embeddings: atom-level *h*_*A*_, fuse-level *h*_*F*_, and mol-level *h*_*M*_. We calculate the similarity *Sim*_*ij*_ of pairs of them.

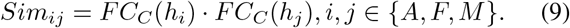

During feature extraction for a set of molecules, if *h*_*i*_ and *h*_*j*_ originate from the same molecule, we use the crossentropy loss function to optimize *Sim*_*ij*_ toward 1; otherwise, it approaches 0. That is, we close the three level representations of same molecular, and keep these away of different molecules.

As for prediction, PyrMol employs a two-layer MLP to obtain prediction scores. We define the objective function for learning as:

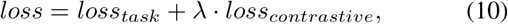

where *λ* is a hyperparameter for hierarchical contrastive learning loss. In experiments, we set the hyperparameter *λ* to 0.2. *loss*_*task*_ is Binary Cross-Entropy Loss for classification tasks and Mean Square Error loss for regression tasks.

## III. EXPERIMENTS AND DISCUSSION

### A. Datasets and Baselines

To evaluate the effectiveness and generalizability of our proposed model, we conducted extensive experiments on ten benchmark datasets from MoleculeNet [27]. These include seven classification tasks—Blood-Brain Barrier Permeability (BBBP), BACE, HIV, ClinTox, Tox21, SIDER, and ToxCast—and three regression tasks—ESOL, FreeSolv, and Lipophilicity. Model performance is primarily assessed using the Area Under the Receiver Operating Characteristic curve (AUC) for classification and Root Mean Square Error (RMSE) for regression. To provide a more granular performance analysis, we also report a comprehensive suite of auxiliary metrics, including Accuracy, Precision, Recall, F1-Score, and AUPR for classification, alongside MSE, MAE, and *R*^2^ for regression. Detailed dataset descriptions are provided in Supplementary Section A.

To demonstrate the competitive advantages of PyrMol, we benchmarked it against 11 state-of-the-art (SOTA) methods. These baselines encompass a diverse range of architectures: traditional graph neural networks (GCN [8], GIN [9], and GAT [10]), specialized molecular graph networks (AttentiveFP [12] and CMPNN [11]), subgraph-enhanced models (HiMol_w/oPretrain [18], PharmHGT [19], and MMGX [21]), and pretraining-based frameworks (HiMol [18], KPGT [28], and MolFCL [17]). Notably, HiMol_w/oPretrain [18] is included as a non-pretrained variant to isolate the architectural contributions of the HiMol framework.

### B. Implementation details

To ensure a rigorous and unbiased evaluation across benchmark datasets, we employ a random scaffold splitting strategy [29]. Unlike the conventional scaffold splitting approach proposed in MoleculeNet [27], which typically sorts scaffolds by frequency and yields deterministic partitions, this strategy introduces stochasticity into the data segmentation. As noted by Deng et al. [29], deterministic splitting procedures can inadvertently favor specific model architectures, thereby inflating perceived generalizability. To mitigate this risk, we eliminate frequency-based sorting and instead shuffle the scaffolds using a random seed. This strategy maintains the structural dissimilarity between training and test sets—crucial for assessing out-of-distribution performance—while providing a more robust partitioning scheme.

All datasets are partitioned into training, validation, and test sets with a ratio of 8:1:1. To achieve statistical significance, we perform 25 independent trials using distinct random seeds. For a fair comparison, all baseline models are implemented via their official repositories using the optimal hyperparameters reported in their respective original studies. Importantly, to ensure an absolutely rigorous and fair comparison and eliminate any bias stemming from data partitioning, all state-of-the-art baseline models were meticulously retrained and evaluated under the exact same random scaffold splitting criterion and random seeds. Our framework is implemented in PyTorch and executed on an NVIDIA Tesla A100 (40GB) GPU, with molecular descriptors (node and edge features) generated using the RDKit library.

### C. Comparison with Baselines

To comprehensively evaluate our framework, we bench-marked PyrMol against 11 state-of-the-art baselines across 10 distinct molecular property datasets. To summarize the global performance, the average ranking (Avg Rank) across all tasks is employed, as presented in Table II. Overall, PyrMol demonstrates commanding superiority, achieving the best Average Rank (1.8) among all 13 models. Specifically, it achieves the top performance on 6 out of 10 tasks, while maintaining highly competitive results on the remaining benchmarks.

**TABLE I.** Node features of each type of node in molecular pyramid graph.

| Node | Dimension | Feature Type |
| --- | --- | --- |
| Atom | 42 | Atom type, atom degree, formal charge, Chiral Tag, the total number of implicit, the explicit bonds connected to the atom, hybridization type. |
| Pharmacophore | 6 | H-bond donor, H-bond acceptor, positive, negative, hydrophobic, aromatic[22]. |
| Functional Group | 115 | Predefined types of edges, rings, functional groups. |
| Retrosynthetic Fragments | 27 | BRICS reaction type features[23]. |
| Molecule | 300 | The average of atoms. |

**TABLE II.** Performance of PyrMol and baseline methods on molecular property prediction.

| Models | Classification(AUC↑) | | | Multi-task Classification(AUC↑) | | | | Regression(RMSE↓) | | | Avg Rank↑ |
| --- | --- | --- | --- | --- | --- | --- | --- | --- | --- | --- | --- |
|  | BBBP | BACE | HIV | ClinTox | SIDER | Tox21 | ToxCast | FreeSolv | Lipophilicity | ESOL |  |
| GCNs | 0.874 | 0.806 | 0.765 | 0.842 | 0.588 | 0.825 | 0.701 | 2.893 | 0.803 | 1.359 | 7.7 |
| [8] | (0.040) | (0.055) | (0.034) | (0.064) | (0.029) | (0.012) | (0.012) | (0.818) | (0.057) | (0.527) |  |
| GINs | 0.881 | 0.829 | 0.761 | 0.835 | 0.596 | 0.824 | 0.698 | 3.046 | 0.764 | 2.165 | 7.7 |
| [9] | (0.040) | (0.058) | (0.044) | (0.070) | (0.029) | (0.013) | (0.015) | (0.842) | (0.041) | (1.438) |  |
| GATs | 0.878 | 0.813 | 0.766 | 0.851 | 0.567 | 0.814 | 0.681 | 2.916 | 0.766 | 1.508 | 8 |
| [10] | (0.037) | (0.048) | (0.037) | (0.058) | (0.023) | (0.012) | (0.014) | (0.739) | (0.051) | (0.596) |  |
| AttentiveFP | 0.860 | 0.820 | 0.673 | 0.746 | 0.526 | 0.768 | 0.627 | 2.274 | 0.963 | 0.893 | 9.4 |
| [12] | (0.591) | (0.037) | (0.041) | (0.111) | (0.025) | (0.016) | (0.018) | (0.367) | (0.103) | (0.119) |  |
| CMPNN | 0.851 | 0.840 | 0.746 | 0.842 | 0.535 | 0.795 | 0.654 | 1.784 | 0.627 | 0.777 | 6.9 |
| [11] | (0.128) | (0.052) | (0.038) | (0.045) | (0.025) | (0.021) | (0.016) | (0.427) | (0.027) | (0.085) |  |
| Himol_w/oPretrain | 0.839 | 0.802 | 0.749 | 0.655 | 0.580 | 0.832 | 0.708 | 3.328 | 0.815 | 1.033 | 8.8 |
| [18] | (0.061) | (0.044) | (0.052) | (0.091) | (0.030) | (0.011) | (0.014) | (1.042) | (0.057) | (0.210) |  |
| PharmHGT | 0.932 | 0.834 | 0.764 | 0.866 | 0.619 | **0.834** | 0.717 | 2.361 | 0.707 | 0.956 | 4.1 |
| [19] | (0.029) | (0.044) | (0.040) | (0.057) | (0.029) | (0.011) | (0.010) | (0.621) | (0.037) | (0.200) |  |
| MMGX | 0.875 | 0.798 | 0.785 | 0.597 | 0.511 | 0.827 | 0.697 | 3.714 | 0.846 | 1.791 | 9.5 |
| [21] | (0.044) | (0.119) | (0.044) | (0.119) | (0.022) | (0.012) | (0.011) | (0.885) | (0.057) | (0.419) |  |
| HiMol | 0.889 | 0.822 | 0.770 | 0.639 | 0.581 | 0.721 | 0.721 | 2.695 | 0.759 | 1.040 | 6.9 |
| [18] | (0.031) | (0.063) | (0.039) | (0.082) | (0.028) | (0.010) | (0.010) | (0.811) | (0.050) | (0.219) |  |
| KPGT | 0.922 | 0.855 | 0.791 | 0.854 | **0.635** | **0.834** | 0.704 | 2.130 | **0.596** | 0.943 | <u>3.2</u> |
| [28] | (0.025) | (0.033) | (0.036) | (0.064) | (0.015) | (0.015) | (0.009) | (0.660) | (0.018) | (0.143) |  |
| MolFCL | 0.872 | 0.833 | 0.796 | 0.862 | 0.598 | 0.808 | 0.711 | 1.777 | 0.638 | **0.769** | 4.2 |
| [17] | (0.070) | (0.046) | (0.037) | (0.077) | (0.033) | (0.020) | (0.016) | (0.494) | (0.029) | (0.091) |  |
| PyrMol | **0.949** | **0.871** | **0.811** | **0.878** | 0.622 | **0.834** | **0.722** | **1.697** | 0.668 | 0.865 | **1.6** |
|  | (0.020) | (0.036) | (0.041) | (0.034) | (0.026) | (0.020) | (0.009) | (0.400) | (0.040) | (0.120) |  |

The bold and underlined values indicate the best and second best results, respectively. ↑ means the higher the value, the better; ↓ means the lower the value, the better.

#### 1) Superiority over Conventional GNNs

When compared to traditional end-to-end graph networks (i.e., GCNs, GINs, GATs) and specialized molecular architectures (AttentiveFP, CMPNN), PyrMol exhibits substantial performance improvements. For instance, on the classification tasks, PyrMol achieves remarkable AUC scores of 0.949 on BBBP, 0.871 on BACE, and 0.811 on HIV. These represent absolute improvements of 6.8%, 3.1%, and 4.5%, respectively, over the strongest baselines in these categories. Furthermore, on the FreeSolv regression task, PyrMol reduces the RMSE to 1.697. These significant improvements empirically validate that explicitly orchestrating molecular topologies into a hierarchical pyramid graph—rather than relying solely on single-scale atomistic topology—profoundly enhances the model’s capacity to capture complex chemical environments.

#### 2) Advantage over Subgraph-Enhanced Models

In contrast to subgraph-enhanced models, PyrMol consistently surpasses single-perspective models like PharmHGT. More importantly, PyrMol decisively outperforms MMGX, a competitive model that also utilizes multiple subgraphs but lacks an adaptive fusion strategy (Avg Rank: 1.8 vs. 10.1). This comparison highlights the critical advantage of our Multi-source Knowledge Enhancement and Fusion Module. Merely extracting diverse substructures is insufficient, and dynamically balancing their complementarity and redundancy through principled heterogeneous message passing is essential for optimal feature extraction.

#### 3) Efficiency against Pretraining Models

Furthermore, we compared PyrMol against state-of-the-art pretraining frameworks (e.g., KPGT, MolFCL, HiMol). Remarkably, PyrMol achieves a superior overall ranking without requiring computationally exhaustive, massive-scale pretraining phases. By strategically synergizing explicit domain priors with simple implicit computational semantics (Mol2Vec), our data-knowledge dual-driven paradigm emerges as a highly efficient alternative to complex pretraining strategies.

Additional evaluation metrics, e.g., Accuracy for classification and MSE for regression, are provided in Supplementary Section B.

### D. Validation of the Pyramid Molecular Graph

To validate the “plug-and-play” versatility of the proposed Pyramid Molecular Graph (PMG) and further demonstrate its intrinsic topological value, we explicitly replaced the native atomic graphs (or pre-defined hierarchical graphs) of five established baselines with our PMG module. The evaluated baselines encompass both classical foundational GNNs (GCNs, GINs, GATs) and advanced subgraph-augmented architectures (HiMol_w/oPretrain, PharmHGT).

We evaluated these PMG-enhanced variants across six representative datasets. As comprehensively visualized in the relative performance gain heatmap (Fig.2 b) and performance comparison bar graph(Fig.2 a). The specific details showed in Supplementary Table S11, the integration of the PMG module yielded performance improvements in 24 out of the 30 experimental evaluations.

**Fig. 2.**
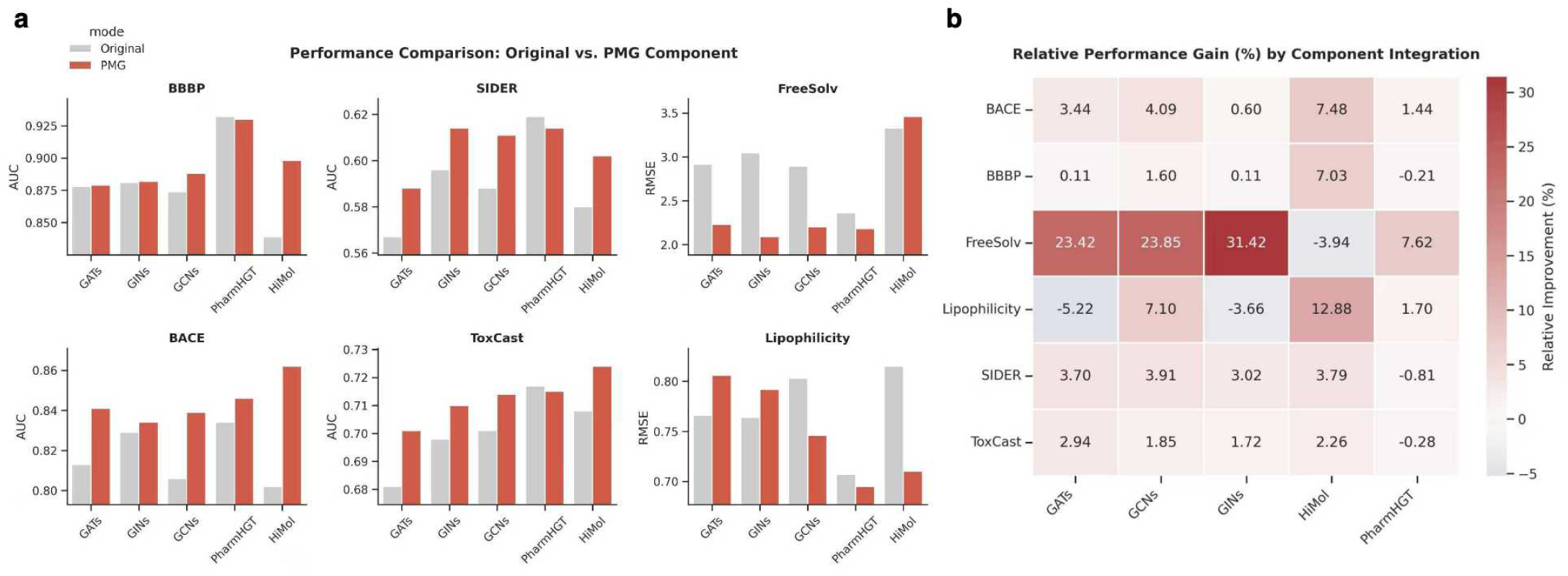
Relative Performance Gain of the baselines with the Pyramid Molecular Graph. a is performance comparison bar graph between original models and the models added PMG Component. b is the relative performance gain heatmap

Notably, foundational architectures that inherently lack explicit domain knowledge experienced the most profound benefits. GCNs, the most fundamental architecture evaluated, achieved consistent gains across all six datasets with an impressive average performance improvement of 7.06%. Similarly, GINs and GATs secured average overall boosts of 5.54% and 4.73%, respectively. The impact of PMG acting as a transformative structural prior is most acutely observed on the FreeSolv dataset, where PMG integration catapulted the performance of GINs, GCNs, and GATs by exceptional margins of 31.42% (i.e., (3.046 − 2.089)/3.046), 23.85%, and 23.42%, respectively. This underscores the capacity of the pyramid topology to empower simple architectures to extract highly expressive, expert-level structural features.

Furthermore, leading subgraph-enhanced baselines also exhibited robust performance enhancements, with the PMG-augmented versions of HiMol_w/oPretrain and PharmHGT achieving average gains of 4.92% and 1.58% over their original implementations.

A particularly compelling observation emerges when evaluating the necessity of massive pretraining. As detailed in Supplementary Table S12, substituting the native topology with our PMG module—without utilizing any pretraining phase (HiMol_PMG)—actually surpasses the fully pretrained HiMol model on five out of six benchmarks (BBBP, BACE, SIDER, ToxCast, and Lipophilicity). For instance, on the BBBP dataset, HiMol_PMG achieves an AUC of 0.898, outperforming both Himol_w/oPretrain (0.839) and the pretrained Himol (0.889). This finding implies that explicitly guiding models with structured, multi-scale domain priors is a highly effective strategy, offering a computationally lightweight complement or alternative to having models implicitly learn these semantics through extensive pretraining.

Collectively, these robust statistical averages and the profound performance shifts against pretrained models validate that our proposed PMG acts as a universal, framework-agnostic enhancement mechanism. By explicitly orchestrating multi-level node message passing, PMG fundamentally elevates the feature learning efficiency of diverse graph neural networks.

### E. Effectiveness Analysis of Multi-source Knowledge Enhancement and Fusion

To rigorously validate the strict necessity of integrating diverse chemical expertise, we conducted an ablation study exploring various combinations of prior knowledge. Since the atomic graph (A) provides the fundamental molecular topology, we established it as the universal base graph. Subsequently, we evaluated seven distinct subgraph-enhanced modes by systematically augmenting the atomic graph with functional groups (F), pharmacophores (P), and retrosynthetic fragments (R). The performance across diverse classification and regression benchmarks is summarized in Table III.

**TABLE III.** Model performance of multiple graph combination models.

| Subgraphs | Classification(AUC↑) | | | | Regression(RMSE↓) | |
| --- | --- | --- | --- | --- | --- | --- |
|  | BBBP | BACE | SIDER | ToxCast | FreeSolv | Lipophilicity |
| A | 0.878 | 0.813 | 0.566 | 0.671 | 2.916 | 0.766 |
|  | (0.037) | (0.048) | (0.025) | (0.014) | (0.739) | (0.051) |
| A+R | 0.909 | 0.852 | 0.553 | 0.695 | 2.874 | 0.703 |
|  | (0.034) | (0.053) | (0.024) | (0.014) | (0.770) | (0.025) |
| A+F | 0.928 | **0.859** | 0.548 | 0.691 | 2.541 | 0.666 |
|  | (0.015) | (0.039) | (0.034) | (0.014) | (0.477) | (0.043) |
| A+P | 0.908 | 0.859 | 0.551 | 0.699 | 2.614 | 0.671 |
|  | (0.033) | (0.043) | (0.041) | (0.007) | (0.555) | (0.025) |
| A+R+F | 0.930 | 0.852 | 0.553 | 0.690 | 2.639 | 0.673 |
|  | (0.026) | (0.032) | (0.031) | (0.018) | (0.565) | (0.017) |
| A+R+P | 0.922 | 0.849 | 0.551 | **0.700** | 2.540 | 0.683 |
|  | (0.032) | (0.034) | (0.027) | (0.013) | (0.582) | (0.027) |
| A+F+P | 0.930 | 0.852 | 0.551 | 0.698 | 2.520 | 0.680 |
|  | (0.020) | (0.038) | (0.040) | (0.014) | (0.439) | (0.038) |
| A+R+F+P | **0.934** | 0.850 | **0.559** | 0.695 | **2.475** | **0.656** |
|  | (0.030) | (0.033) | (0.024) | (0.024) | (0.534) | (0.022) |

The bold values indicate the best results. A, R, F, P means atomic graphs, retrosynthetic fragments, functional groups, and pharmacophores, respectively.

Evaluating the single-knowledge enhancements (A+R, A+F, and A+P) reveals a clear task-dependent preference. For instance, on the BBBP dataset, incorporating functional groups (A+F) yields a significantly better AUC (0.928) compared to pharmacophores (A+P, 0.908) or reaction rules (A+R, 0.909). Conversely, A+P and A+R demonstrate unique advantages on multi-classification tasks. This observation perfectly aligns with our core hypothesis: distinct molecular properties are governed by varying physical and chemical mechanisms. Consequently, relying on any single-perspective subgraph is inherently insufficient for universal generalization across diverse biochemical tasks.

Furthermore, as progressively more knowledge sources are integrated, the predictive performance consistently improves, eventually reaching the optimal performance of the fully integrated A+R+F+P model. As shown in Table III, the full combination achieves the highest predictive capability across almost all metrics, including the lowest RMSE on FreeSolv (2.475) and Lipophilicity (0.656), as well as the highest AUC on BBBP (0.934) and SIDER (0.559).

These results underscore that the integration of functional groups (F), pharmacophores (P), and retrosynthetic fragments (R) priors is not merely additive, but highly synergistic. Equipping the model with this comprehensive, complementary knowledge base is essential

### F. Ablation Study

To rigorously evaluate the contribution of each component in PyrMol, we conducted a comprehensive ablation study across six benchmark datasets. We designed five variants to analyze the specific effects of the Pyramid Molecular Graph (PMG), Hierarchical Contrastive Learning Strategy (CL), Multi-source Knowledge Enhancement and Fusion Module (EF), and the integration of Mol2Vec features (M2V). The results are shown in Fig.3.

**Fig. 3.**
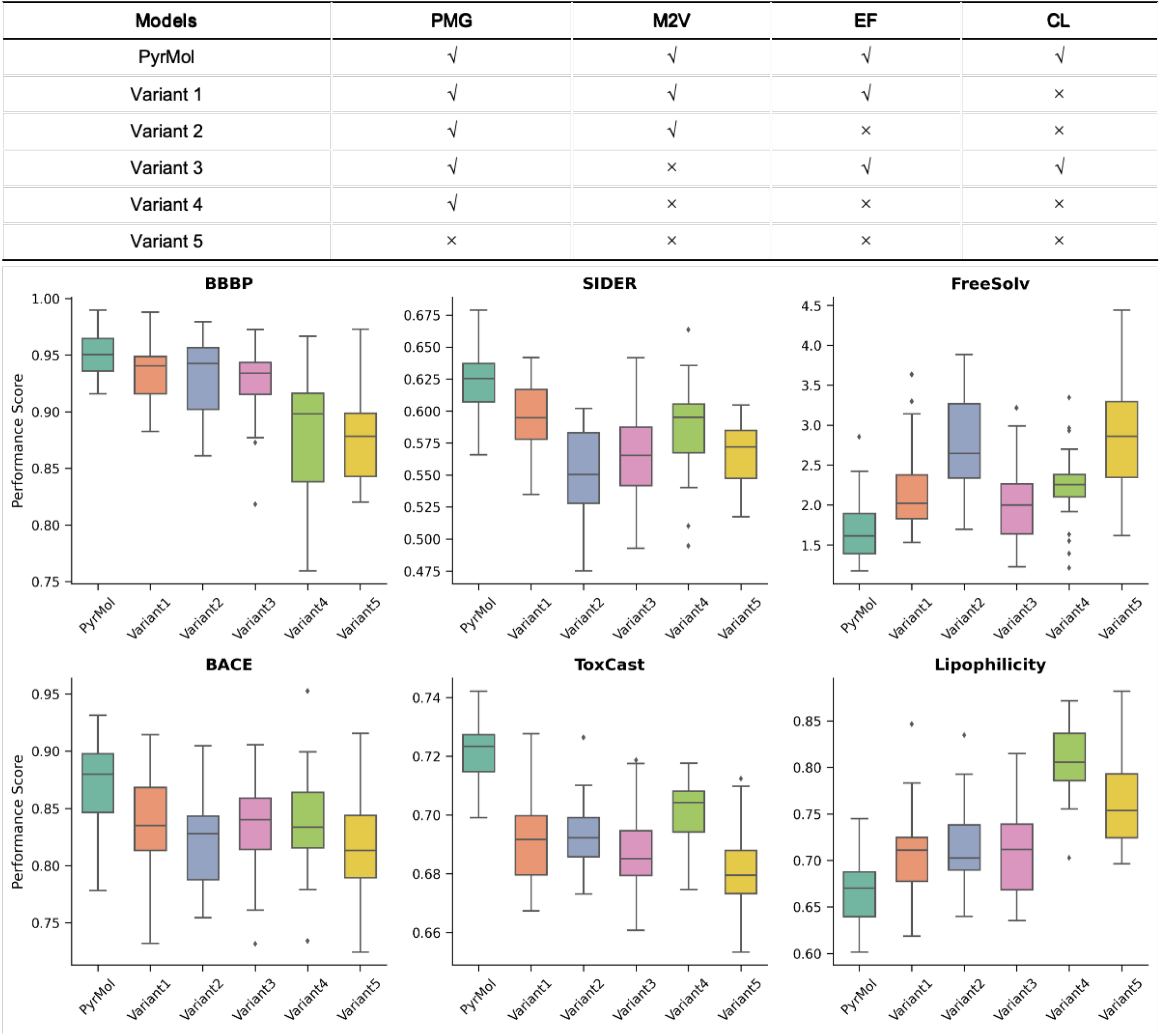
The results of ablation study on six datasets. The figure contains models definition details introduction and performance boxplot figure.

Comparing Variant 4 with the Variant 5 demonstrates the intrinsic superiority of our topological design. Even without auxiliary features or learning strategies, the Pyramid Hierarchical Graph construction significantly outperforms the Variant 5, particularly in classification tasks. Besides, on the FreeSolv dataset, Variant 4 reduces the RMSE from 2.916 to 2.233. This indicates that our Pyramid Hierarchical Graph construction effectively captures multi-scale structural information inherent in molecular graphs, providing a robust geometric foundation.

Notably, the complete PyrMol framework substantially out-performs both Variant 2 and Variant 3. The significant performance drop observed in both partial configurations indicates a powerful synergistic effect. It demonstrates that simply supplementing initial node features via Mol2Vec is insufficient; rather, it provides a foundational semantic space that strictly requires our EF and CL modules to be effectively aligned, refined, and translated into superior predictive performance.

The comparison between Variant 1 and PyrMol highlights that the CL module consistently refines the feature space across all six tasks, yielding universal performance improvements (e.g., pushing the AUC on BBBP from 0.936 to 0.949). Finally, the full PyrMol model achieves the best performance across all metrics. While Mol2Vec provides a useful initialization, the dominant performance gains are driven by the synergy of our Pyramid architecture and learning strategies, confirming that our proposed modules are the primary drivers of PyrMol’s performance.

### G. Interpretation Results

A key advantage of PyrMol is its inherent interpretability. Because PyrMol utilizes the GAT architecture to aggregate features from substructure nodes to the global molecular node, and the hierarchical mapping between atoms and substructures is deterministically defined within the pyramid graph, the model can explicitly trace and project learned attention scores back to individual atoms. To validate this capability, we visualize the atomic attention weights for representative molecules from the BBBP dataset. According to the visualization scheme, dark red indicates the highest attention score, transitioning through light red to light blue, with dark blue representing the lowest weight.

As illustrated in Fig. 4A, the first molecule ([*C*@@*H*]3(*C*1 = *CC* = *CC* = *C*1)*CN*2*CCSC*2 = *N* 3) features a rigid planar structure formed by two fused heterocycles attached to a benzene ring. Chemically, the sulfur atom in the thiazole ring exhibits lower polarity and stronger hydrophobicity than typical oxygen analogues, thereby positively contributing to BBB permeability. Crucially, PyrMol captures this phenomenon, assigning the highest attention weight (visualized as dark red) to the sulfur atom. Conversely, the *sp*^2^ hybridized nitrogen atom acts as a strong hydrogen bond acceptor, typically impeding BBB penetration. The model accurately identifies this resistance point, assigning it a negatively correlated, lower attention score (visualized as blue).

**Fig. 4.**
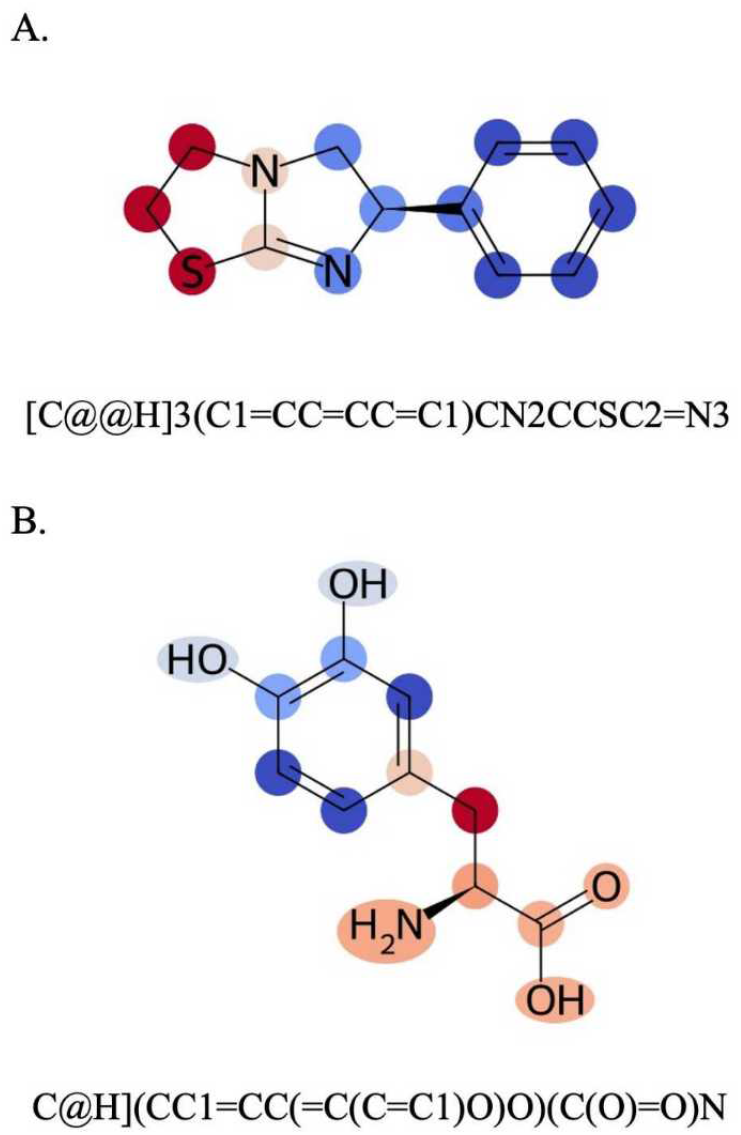
Interpretability analysis of attention scores on the BBBP dataset. Dark red indicates the atom have the highest attention score, light red the next highest, and as the score decreases, it changes from light red to light blue, with dark blue representing the lowest weight.

As shown in Fig. 4B, the second molecule (*C*[@*H*](*CC*1 = *CC*(= *C*(*C* = *C*1)*O*)*O*)(*C*(*O*) = *O*)*N*) comprises an amino acid backbone, an aromatic side chain, catechol hydroxyl groups, and a methylene bridge. PyrMol correctly recognizes the amino acid backbone (*H*_2_*N* − *CH* − *COOH*) with a high correlation for BBB crossing (visualized as light red, indicating the next highest attention). Meanwhile, the crucial hydrophobic methylene bridge (−*CH*_2_−) connecting the main and side chains is rightly highlighted with the highest attention score (dark red).

Collectively, these visualizations confirm that the attention mechanisms within PyrMol align with sophisticated pharmaceutical intuition. By focusing on atoms and substructures that dictate macroscopic properties, PyrMol provides highly reliable and explainable clues for downstream molecular optimization.

## IV. Conclusion

In this work, we propose PyrMol, a knowledge-structured heterogeneous pyramid framework that pioneers a data-knowledge dual-driven paradigm for molecular representation learning. Moving beyond conventional single-scale atomic topology, PyrMol systematically mirrors the cognitive hierarchy of expert chemists. Its core pyramid graph orchestrates message passing across multi-scale chemical environments, while the adaptive Multi-source Knowledge Enhancement and Fusion module dynamically balances the complementarity and redundancy of heterogeneous domain priors. Complemented by a hierarchical contrastive learning strategy, PyrMol significantly amplifies the discriminative power of molecular embeddings. Extensive evaluations across ten benchmarks demonstrate that PyrMol consistently outperforms twelve state-of-the-art baselines. Remarkably, without relying on computationally exhaustive pretraining phases, our end-to-end framework establishes a highly efficient and robust reference paradigm, proving the immense potential of explicitly integrating multi-perspective structural knowledge.

While PyrMol demonstrates competitive performance on 2D molecular graphs, there are several directions for future improvement. Currently, the framework integrates three specific types of domain knowledge; however, incorporating a broader range of reliable features could further improve molecular representations. Furthermore, extending the current pyramid structure to encompass three-dimensional (3D) molecular conformations is a necessary next step to better capture stereochemical structure-property relationships. We hope that PyrMol can serve as a practical and effective framework for molecular representation learning, providing useful computational support for AI-aided drug discovery.

## Acknowledgments

This work was supported in part by the National Natural Science Foundation of China (Grant Nos. U25A20449, 62272309 and U22A2041). This work was carried out in part using computing resources at the High-Performance Computing Center of Central South University.

