## Supplemental Tables 1-11 for "PyrMol: A Knowledge-Structured Pyramid Graph Framework for Generalizable Molecular Property Prediction"

#### A Datasets description

##### A.1 Binary classification tasks

The BBBP dataset comprises 2,035 molecules annotated with binary labels indicating blood-brain barrier permeability, making it a benchmark for predicting whether compounds can cross the blood-brain barrier. The BACE dataset includes 1,513 molecules and offers both quantitative ($IC_{50}$) and qualitative (binary) binding measurements for inhibitors of human $\beta$-secretase 1 (BACE-1). The HIV dataset contains 41,127 compounds screened for their ability to inhibit HIV replication.

##### A.2 Multi-class classification tasks

The ClinTox dataset is curated from 1,468 molecules, offering a binary classification task to distinguish FDA-approved drugs from those that failed in clinical trials owing to toxicity. The Tox21 dataset, originating from the Tox21 Challenge, provides assay results for 7,821 molecules across 12 distinct targets related to drug-induced toxicity. For the SIDER dataset, it encompasses 1,379 approved drugs, with side effects annotated and organized into 27 system organ classes. We also include the ToxCast dataset, which aggregates in vitro high-throughput screening data for 8,597 compounds over 617 toxicological tasks.

##### A.3 Regression tasks

For predicting key physicochemical properties, we employ three benchmarks. The ESOL dataset offers aqueous solubility measurements for 1,128 compounds. FreeSolv is a curated set of 642 molecules with corresponding experimental hydration free energies. The Lipophilicity dataset includes 4,198 compounds annotated with their experimental octanol/water distribution coefficients (logD at pH 7.4).

#### B Performance of PyrMol and baseline methods

Additional evaluation metrics, e.g., Accuracy, Precision, Recall, F1-Score, AUPR and AUC for classification and MSE,RMSE,MAE,R2 for regression, are provided in Table S1-S10.

**Table S1.** Performance of PyrMol and baseline methods on the BBBP dataset.

| Models | Accuracy | Precision | Recall | F1-Score | AUPR | AUC |
|---|---|---|---|---|---|---|
| GCNs | 0.8571(0.0350) | 0.9047(0.0296) | 0.9118(0.0565) | 0.9066(0.0270) | 0.9478(0.0301) | 0.8742(0.0409) |
| GINs | 0.8613(0.0359) | 0.9007(0.0328) | 0.9216(0.0470) | 0.9099(0.0261) | 0.9457(0.0362) | 0.8814(0.0406) |
| GATs | 0.8579(0.0364) | 0.9041(0.0374) | 0.9121(0.0433) | 0.9071(0.0271) | 0.9480(0.0314) | 0.8786(0.0376) |
| AttentiveFP | 0.8672(0.0285) | 0.8897(0.0365) | 0.9455(0.0322) | 0.9158(0.0191) | 0.9413(0.0236) | 0.8600(0.0592) |
| CMPNN | 0.8219(0.0888) | 0.9060(0.0516) | 0.8593(0.0938) | 0.8792(0.0625) | 0.9445(0.0434) | 0.8518(0.1286) |
| Himol_w/oPretrain | 0.8404(0.0426) | 0.8922(0.0320) | 0.9027(0.0668) | 0.8954(0.0340) | 0.9357(0.0298) | 0.8391(0.0615) |
| PharmHGT | 0.8888(0.0322) | 0.9204(0.0403) | 0.9377(0.0440) | 0.9276(0.0234) | 0.9775(0.0134) | 0.9324(0.0295) |
| MMGX | 0.8143(0.0481) | 0.9207(0.0321) | 0.8295(0.0712) | 0.8708(0.0424) | 0.9518(0.0213) | 0.8750(0.0446) |
| HiMol | 0.8731(0.0291) | 0.9131(0.0290) | 0.9220(0.0354) | 0.9169(0.0220) | 0.9577(0.0164) | 0.8893(0.0319) |
| KPGT | 0.8928(0.0299) | 0.9087(0.0336) | 0.9543(0.0232) | 0.9306(0.0227) | 0.9734(0.0097) | 0.9223(0.0257) |
| MolFCL | 0.8264(0.0861) | 0.9097(0.0406) | 0.8629(0.1220) | 0.8798(0.0695) | 0.9484(0.0333) | 0.8723(0.0706) |
| PyrMol | 0.8988(0.0324) | 0.9358(0.0299) | 0.9318(0.0438) | 0.9329(0.0243) | 0.9831(0.0094) | 0.9490(0.0200) |

**Table S2.** Performance of PyrMol and baseline methods on the BACE dataset.

| Models | Accuracy | Precision | Recall | F1-Score | AUPR | AUC |
|---|---|---|---|---|---|---|
| GCNs | 0.7316(0.0519) | 0.6976(0.0866) | 0.7731(0.1461) | 0.7217(0.0935) | 0.7840(0.0872) | 0.8066(0.0555) |
| GINs | 0.7550(0.0555) | 0.7245(0.0763) | 0.7956(0.1325) | 0.7497(0.0714) | 0.8023(0.0839) | 0.8294(0.0580) |
| GATs | 0.7431(0.0503) | 0.6986(0.0967) | 0.8042(0.1102) | 0.7409(0.0793) | 0.7835(0.0873) | 0.8130(0.0483) |
| AttentiveFP | 0.7311(0.0893) | 0.7467(0.0975) | 0.6647(0.1807) | 0.6783(0.1571) | 0.7679(0.0823) | 0.8204(0.0377) |
| CMPNN | 0.7605(0.1091) | 0.7542(0.1242) | 0.7659(0.1297) | 0.7454(0.0960) | 0.7875(0.1000) | 0.8403(0.0522) |
| Himol_w/oPretrain | 0.7200(0.0696) | 0.7511(0.0595) | 0.6043(0.2017) | 0.6469(0.1279) | 0.7541(0.0724) | 0.8023(0.0443) |
| PharmHGT | 0.7734(0.0469) | 0.7674(0.0674) | 0.7601(0.1110) | 0.7573(0.0630) | 0.8085(0.0715) | 0.8341(0.0449) |
| MMGX | 0.7348(0.1198) | 0.7467(0.1333) | 0.7317(0.1918) | 0.7065(0.1639) | 0.7620(0.1285) | 0.7984(0.1197) |
| HiMol | 0.7574(0.0612) | 0.7488(0.0773) | 0.7030(0.1534) | 0.7164(0.1033) | 0.7781(0.0909) | 0.8224(0.0632) |
| KPGT | 0.8559(0.0370) | 0.7722(0.0752) | 0.7767(0.1118) | 0.7678(0.0646) | 0.8171(0.0667) | 0.8559(0.0337) |
| MolFCL | 0.7666(0.0552) | 0.7405(0.0839) | 0.7636(0.1468) | 0.7425(0.0889) | 0.7852(0.0836) | 0.8330(0.0464) |
| PyrMol | 0.7939(0.0387) | 0.7509(0.0733) | 0.8350(0.0904) | 0.7857(0.0537) | 0.8331(0.0680) | 0.8715(0.0369) |

**Table S3.** Performance of PyrMol and baseline methods on the HIV dataset.

| Models | Accuracy | Precision | Recall | F1-Score | AUPR | AUC |
|---|---|---|---|---|---|---|
| GCNs | 0.9642(0.0055) | 0.5544(0.1384) | 0.2680(0.0972) | 0.3427(0.0965) | 0.3209(0.0788) | 0.7652(0.0348) |
| GINs | 0.9663(0.0050) | 0.5841(0.0936) | 0.2744(0.0889) | 0.3650(0.0872) | 0.3368(0.0927) | 0.7616(0.0447) |
| GATs | 0.9641(0.0060) | 0.5221(0.0964) | 0.3003(0.0899) | 0.3727(0.0821) | 0.3261(0.0799) | 0.7662(0.0375) |
| AttentiveFP | 0.9635(0.0073) | 0.4372(0.3834) | 0.0197(0.0233) | 0.0365(0.0418) | 0.1692(0.0397) | 0.6734(0.0418) |
| CMPNN | 0.9552(0.0380) | 0.5364(0.1886) | 0.2270(0.1009) | 0.2872(0.1029) | 0.2764(0.0813) | 0.7461(0.0383) |
| Himol_w/oPretrain | 0.9648(0.0074) | 0.5816(0.2219) | 0.1525(0.0876) | 0.2283(0.1161) | 0.2949(0.0956) | 0.7494(0.0525) |
| PharmHGT | 0.9659(0.0053) | 0.5793(0.1144) | 0.2553(0.0947) | 0.3430(0.1109) | 0.3318(0.0863) | 0.7649(0.0406) |
| MMGX | 0.8192(0.0840) | 0.1363(0.0528) | 0.6146(0.1139) | 0.2166(0.0661) | 0.3200(0.0684) | 0.7850(0.0448) |
| HiMol | 0.9661(0.0066) | 0.6337(0.1679) | 0.1944(0.0882) | 0.2843(0.0989) | 0.3373(0.0721) | 0.7709(0.0396) |
| KPGT | 0.9682(0.0057) | 0.6656(0.0867) | 0.2828(0.0650) | 0.3902(0.0645) | 0.3887(0.0692) | 0.7910(0.0360) |
| MolFCL | 0.9659(0.0067) | 0.5810(0.1233) | 0.3078(0.0661) | 0.3957(0.0727) | 0.3702(0.0851) | 0.7965(0.0374) |
| PyrMol | 0.9662(0.0059) | 0.5942(0.1253) | 0.2809(0.1043) | 0.3672(0.1057) | 0.3695(0.0877) | 0.8108(0.0414) |

**Table S4.** Performance of PyrMol and baseline methods on the ClinTox dataset.

| Models | Accuracy | Precision | Recall | F1-Score | AUPR | AUC |
|---|---|---|---|---|---|---|
| GCNs | 0.9103(0.0259) | 0.6505(0.1164) | 0.6461(0.1043) | 0.6429(0.1057) | 0.7060(0.0647) | 0.8421(0.0644) |
| GINs | 0.9077(0.0355) | 0.6987(0.1400) | 0.6549(0.1017) | 0.6568(0.0944) | 0.7068(0.0663) | 0.8357(0.0700) |
| GATs | 0.9064(0.0256) | 0.6857(0.0925) | 0.6611(0.0866) | 0.6620(0.0787) | 0.7126(0.0588) | 0.8519(0.0581) |
| AttentiveFP | 0.9166(0.0211) | 0.4938(0.1105) | 0.5131(0.0512) | 0.4984(0.0689) | 0.6239(0.0798) | 0.7466(0.1111) |
| CMPNN | 0.7995(0.0835) | 0.6266(0.0520) | 0.7647(0.0745) | 0.6359(0.0623) | 0.6710(0.0644) | 0.8424(0.0459) |
| Himol_w/oPretrain | 0.1710(0.0609) | 0.5439(0.0201) | 0.5312(0.0440) | 0.1733(0.0598) | 0.5759(0.0403) | 0.6553(0.0914) |
| PharmHGT | 0.9261(0.0243) | 0.7539(0.1086) | 0.6641(0.0860) | 0.6866(0.0865) | 0.7549(0.0584) | 0.8664(0.0572) |
| MMGX | 0.4083(0.2434) | 0.5363(0.0367) | 0.4555(0.1764) | 0.2722(0.1852) | 0.5621(0.0504) | 0.5974(0.1194) |
| HiMol | 0.1693(0.0601) | 0.5372(0.0192) | 0.5164(0.0556) | 0.1656(0.0606) | 0.5787(0.0415) | 0.6399(0.0823) |
| KPGT | 0.9214(0.0217) | 0.7574(0.1846) | 0.5705(0.0568) | 0.5949(0.0805) | 0.7313(0.0618) | 0.8543(0.0646) |
| MolFCL | 0.8046(0.1430) | 0.6795(0.1008) | 0.7634(0.0904) | 0.6563(0.1103) | 0.7180(0.0767) | 0.8626(0.0774) |
| PyrMol | 0.9163(0.0235) | 0.6849(0.1613) | 0.6396(0.1085) | 0.6399(0.1092) | 0.7399(0.0646) | 0.8784(0.0342) |

**Table S5.** Performance of PyrMol and baseline methods on the SIDER dataset.

| Models | Accuracy | Precision | Recall | F1-Score | AUPR | AUC |
|---|---|---|---|---|---|---|
| GCNs | 0.7621(0.0152) | 0.5783(0.0405) | 0.6660(0.0274) | 0.6073(0.0235) | 0.6548(0.0248) | 0.5888(0.0296) |
| GINs | 0.7581(0.0180) | 0.5927(0.0371) | 0.6526(0.0417) | 0.6096(0.0286) | 0.6603(0.0246) | 0.5963(0.0294) |
| GATs | 0.7609(0.0134) | 0.5607(0.0264) | 0.6746(0.0255) | 0.6039(0.0207) | 0.6417(0.0221) | 0.5678(0.0238) |
| AttentiveFP | 0.7562(0.0190) | 0.5219(0.0287) | 0.6733(0.0447) | 0.5812(0.0346) | 0.6126(0.0247) | 0.5263(0.0257) |
| CMPNN | 0.6258(0.0620) | 0.6009(0.0206) | 0.6410(0.0909) | 0.5924(0.0504) | 0.6204(0.0246) | 0.5356(0.0250) |
| Himol_w/oPretrain | 0.4920(0.0626) | 0.6167(0.0256) | 0.6661(0.1034) | 0.5311(0.0880) | 0.6458(0.0218) | 0.5806(0.0308) |
| PharmHGT | 0.7501(0.0219) | 0.6256(0.0317) | 0.6598(0.0476) | 0.6298(0.0272) | 0.6745(0.0241) | 0.6196(0.0292) |
| MMGX | 0.4413(0.0613) | 0.5820(0.0559) | 0.5040(0.0754) | 0.3449(0.0570) | 0.6085(0.0258) | 0.5110(0.0228) |
| HiMol | 0.4753(0.0710) | 0.6256(0.0371) | 0.6081(0.1817) | 0.4858(0.1393) | 0.6492(0.0280) | 0.5810(0.0283) |
| KPGT | 0.7549(0.0168) | 0.6521(0.0349) | 0.6168(0.0433) | 0.6147(0.0314) | 0.6912(0.0175) | 0.6354(0.0150) |
| MolFCL | 0.6783(0.0383) | 0.6194(0.0263) | 0.7698(0.0773) | 0.6658(0.0315) | 0.6571(0.0281) | 0.5984(0.0337) |
| PyrMol | 0.7543(0.0159) | 0.6127(0.0432) | 0.6489(0.0343) | 0.6220(0.0292) | 0.6702(0.0232) | 0.6223(0.0260) |

**Table S6.** Performance of PyrMol and baseline methods on the Tox21 dataset.

| Models | Accuracy | Precision | Recall | F1-Score | AUPR | AUC |
|---|---|---|---|---|---|---|
| GCNs | 0.9287(0.0042) | 0.6008(0.0621) | 0.3289(0.0520) | 0.4058(0.0482) | 0.4520(0.0322) | 0.8254(0.0120) |
| GINs | 0.9284(0.0049) | 0.6091(0.0570) | 0.3084(0.0520) | 0.3884(0.0480) | 0.4487(0.0383) | 0.8243(0.0138) |
| GATs | 0.9262(0.0046) | 0.5702(0.0621) | 0.2939(0.0395) | 0.3665(0.0378) | 0.4158(0.0312) | 0.8140(0.0128) |
| AttentiveFP | 0.9324(0.0041) | 0.2898(0.1356) | 0.1232(0.0461) | 0.1519(0.0541) | 0.2606(0.0354) | 0.7680(0.0169) |
| CMPNN | 0.8928(0.0175) | 0.3990(0.0626) | 0.3529(0.0711) | 0.3420(0.0352) | 0.3395(0.0368) | 0.7950(0.0219) |
| Himol_w/oPretrain | 0.6237(0.1503) | 0.3449(0.1087) | 0.6821(0.1324) | 0.2868(0.0613) | 0.4386(0.0358) | 0.8325(0.0118) |
| PharmHGT | 0.9294(0.0050) | 0.6142(0.0709) | 0.3478(0.0417) | 0.4277(0.0408) | 0.4773(0.0361) | 0.8343(0.0113) |
| MMGX | 0.7823(0.0283) | 0.2326(0.0252) | 0.7266(0.0384) | 0.3340(0.0244) | 0.3991(0.0299) | 0.8273(0.0125) |
| HiMol | 0.5702(0.1784) | 0.1995(0.0928) | 0.6859(0.1031) | 0.2721(0.0672) | 0.4362(0.0355) | 0.7211(0.0106) |
| KPGT | 0.9171(0.0078) | 0.6842(0.0724) | 0.2353(0.0463) | 0.3252(0.0554) | 0.4836(0.0411) | 0.8343(0.0152) |
| MolFCL | 0.8958(0.0166) | 0.4237(0.0597) | 0.3911(0.0663) | 0.3717(0.0385) | 0.3673(0.0385) | 0.8084(0.0208) |
| PyrMol | 0.7936(0.0397) | 0.2381(0.0319) | 0.7258(0.0688) | 0.3452(0.0355) | 0.4388(0.0467) | 0.8344(0.0206) |

**Table S7.** Performance of PyrMol and baseline methods on the ToxCast dataset.

| Models | Accuracy | Precision | Recall | F1-Score | AUPR | AUC |
|---|---|---|---|---|---|---|
| GCNs | 0.8384(0.0077) | 0.3830(0.0200) | 0.2465(0.0217) | 0.2717(0.0218) | 0.4019(0.0149) | 0.7010(0.0124) |
| GINs | 0.8383(0.0092) | 0.3602(0.0247) | 0.2346(0.0274) | 0.2574(0.0266) | 0.3951(0.0189) | 0.6987(0.0150) |
| GATs | 0.8326(0.0090) | 0.3165(0.0328) | 0.2157(0.0212) | 0.2299(0.0215) | 0.3700(0.0187) | 0.6813(0.0146) |
| AttentiveFP | 0.8627(0.0174) | 0.0068(0.0051) | 0.1341(0.0214) | 0.0085(0.0031) | 0.0512(0.0069) | 0.6275(0.0188) |
| CMPNN | 0.6141(0.0426) | 0.2552(0.0123) | 0.6294(0.0604) | 0.3176(0.0155) | 0.3355(0.0162) | 0.6541(0.0164) |
| Himol_w/oPretrain | 0.5415(0.0744) | 0.3150(0.0451) | 0.6934(0.0788) | 0.3127(0.0348) | 0.4044(0.0206) | 0.7085(0.0146) |
| PharmHGT | 0.8416(0.0070) | 0.4020(0.0220) | 0.2662(0.0218) | 0.2939(0.0214) | 0.4169(0.0152) | 0.7178(0.0106) |
| MMGX | 0.6198(0.0198) | 0.2827(0.0141) | 0.6938(0.0388) | 0.3410(0.0145) | 0.3778(0.0169) | 0.6975(0.0117) |
| HiMol | 0.5268(0.0770) | 0.3311(0.0492) | 0.7010(0.0796) | 0.3072(0.0336) | 0.4189(0.0151) | 0.7211(0.0106) |
| KPGT | 0.8261(0.0078) | 0.3684(0.0351) | 0.1891(0.0195) | 0.2277(0.0223) | 0.4130(0.0216) | 0.7045(0.0095) |
| MolFCL | 0.7634(0.0319) | 0.3614(0.0325) | 0.4593(0.0657) | 0.3560(0.0189) | 0.3975(0.0186) | 0.7114(0.0165) |
| PyrMol | 0.6991(0.0258) | 0.3122(0.0175) | 0.6272(0.0475) | 0.3623(0.0162) | 0.4115(0.0164) | 0.7215(0.0100) |

**Table S8.** Performance of PyrMol and baseline methods on the FreeSolv dataset.

| Models | MSE | RMSE | MAE | R2 |
|---|---|---|---|---|
| GCNs | 9.0412(4.9967) | 2.8933(0.8186) | 2.1160(0.6347) | 0.4760(0.2620) |
| GINs | 9.9877(5.3362) | 3.0461(0.8422) | 2.0752(0.5009) | 0.4335(0.2243) |
| GATs | 9.0513(4.5795) | 2.9163(0.7394) | 2.0771(0.4987) | 0.4909(0.1545) |
| AttentiveFP | 5.3098(1.7631) | 2.2748(0.3673) | 1.6911(0.1782) | 0.6627(0.1554) |
| CMPNN | 3.3672(1.6594) | 1.7845(0.4276) | 1.2806(0.2729) | 0.8064(0.0627) |
| Himol_w/oPretrain | 12.1666(7.2570) | 3.3287(1.0424) | 2.3206(0.7788) | 0.3141(0.3510) |
| PharmHGT | 5.9616(3.1249) | 2.3611(0.6219) | 1.6910(0.4307) | 0.6512(0.1517) |
| MMGX | 14.5832(7.1301) | 3.7148(0.8850) | 2.8834(0.6950) | 0.1443(0.3518) |
| HiMol | 7.9240(4.7931) | 2.6953(0.8119) | 1.8399(0.5635) | 0.5533(0.2241) |
| KPGT | 4.9584(3.4402) | 2.1305(0.6608) | 1.3601(0.3926) | 0.7271(0.1247) |
| MolFCL | 3.4023(1.9643) | 1.7770(0.4947) | 1.2846(0.2823) | 0.8029(0.0791) |
| PyrMol | 3.0423(1.5491) | 1.6977(0.4001) | 1.2079(0.2358) | 0.8259(0.0494) |

**Table S9.** Performance of PyrMol and baseline methods on the Lipophilicity dataset.

| Models | MSE | RMSE | MAE | R2 |
|---|---|---|---|---|
| GCNs | 0.6494(0.0927) | 0.8038(0.0572) | 0.6132(0.0477) | 0.5612(0.0802) |
| GINs | 0.5865(0.0634) | 0.7647(0.0412) | 0.5839(0.0331) | 0.6065(0.0420) |
| GATs | 0.5899(0.0799) | 0.7663(0.0511) | 0.5843(0.0376) | 0.6050(0.0450) |
| AttentiveFP | 0.9396(0.1977) | 0.9638(0.1039) | 0.7686(0.0903) | 0.3655(0.1246) |
| CMPNN | 0.3949(0.0347) | 0.6278(0.0276) | 0.4756(0.0179) | 0.7305(0.0377) |
| Himol_w/oPretrain | 0.6679(0.0951) | 0.8153(0.0575) | 0.6298(0.0481) | 0.5465(0.0671) |
| PharmHGT | 0.5022(0.0537) | 0.7077(0.0377) | 0.5324(0.0290) | 0.6613(0.0498) |
| MMGX | 0.7199(0.1001) | 0.8465(0.0577) | 0.6598(0.0512) | 0.5074(0.0920) |
| HiMol | 0.5788(0.0782) | 0.7591(0.0505) | 0.5749(0.0384) | 0.6074(0.0530) |
| KPGT | 0.3557(0.0225) | 0.5961(0.0188) | 0.4410(0.0170) | 0.7584(0.0315) |
| MolFCL | 0.4089(0.0380) | 0.6388(0.0291) | 0.4825(0.0223) | 0.7222(0.0299) |
| PyrMol | 0.4480(0.0550) | 0.6681(0.0407) | 0.5015(0.0284) | 0.6959(0.0383) |

**Table S10.** Performance of PyrMol and baseline methods on the ESOL dataset.

| Models | MSE | RMSE | MAE | R2 |
|---|---|---|---|---|
| GCNs | 2.1277(1.8537) | 1.3598(0.5279) | 1.1237(0.5008) | 0.4348(0.4685) |
| GINs | 6.7590(9.8025) | 2.1655(1.4386) | 1.7039(1.1561) | 0.4226(1.6496) |
| GATs | 2.6309(2.4135) | 1.5083(0.5965) | 1.2205(0.5508) | 0.3098(0.5614) |
| AttentiveFP | 0.8117(0.2214) | 0.8930(0.1194) | 0.7017(0.0986) | 0.8013(0.0680) |
| CMPNN | 0.6119(0.1350) | 0.7776(0.0851) | 0.6149(0.0755) | 0.8501(0.0454) |
| Himol_w/oPretrain | 1.1126(0.5168) | 1.0335(0.2107) | 0.8268(0.1813) | 0.7361(0.1075) |
| PharmHGT | 0.9553(0.4071) | 0.9565(0.2009) | 0.7420(0.1633) | 0.7394(0.1555) |
| MMGX | 3.3870(1.6167) | 1.7919(0.4195) | 1.5267(0.4220) | 0.1587(0.5303) |
| HiMol | 1.1308(0.4820) | 1.0405(0.2193) | 0.8165(0.1657) | 0.7387(0.0825) |
| KPGT | 0.9099(0.2836) | 0.9435(0.1433) | 0.7313(0.1122) | 0.7957(0.0457) |
| MolFCL | 0.6005(0.1446) | 0.7696(0.0910) | 0.6103(0.0780) | 0.8530(0.0468) |
| PyrMol | 0.7628(0.2417) | 0.8650(0.1207) | 0.6752(0.0933) | 0.8093(0.0817) |

### C Pyramid Molecular Graph

**Table S11.** Performance of the baselines with the Pyramid Molecular Graph.

| Models | Classification(AUC↑) | | | | Regression(RMSE↓) | |
|---|---|---|---|---|---|---|
| | BBBP | BACE | SIDER | ToxCast | FreeSolv | Lipophilicity |
| GATs | 0.878<br>(0.037) | 0.813<br>(0.048) | 0.567<br>(0.023) | 0.681<br>(0.014) | 2.916<br>(0.739) | **0.766**<br>(0.051) |
| GATs_PMG | **0.879**<br>(0.053) | **0.841**<br>(0.044) | **0.588**<br>(0.037) | **0.701**<br>(0.011) | **2.233**<br>(0.460) | 0.806<br>(0.037) |
| GINs | 0.881<br>(0.040) | 0.829<br>(0.058) | 0.596<br>(0.029) | 0.698<br>(0.015) | 3.046<br>(0.842) | **0.764**<br>(0.041) |
| GINs_PMG | **0.882**<br>(0.056) | **0.834**<br>(0.048) | **0.614**<br>(0.031) | **0.710**<br>(0.010) | **2.089**<br>(0.459) | 0.792<br>(0.040) |
| GCNs | 0.874<br>(0.040) | 0.806<br>(0.055) | 0.588<br>(0.029) | 0.701<br>(0.012) | 2.893<br>(0.818) | 0.803<br>(0.057) |
| GCNs_PMG | **0.888**<br>(0.044) | **0.839**<br>(0.042) | **0.611**<br>(0.030) | **0.714**<br>(0.013) | **2.203**<br>(0.544) | **0.746**<br>(0.033) |
| PharmHGT | 0.932<br>(0.029) | 0.834<br>(0.044) | **0.619**<br>(0.029) | **0.717**<br>(0.010) | 2.361<br>(0.621) | 0.707<br>(0.037) |
| PharmHGT<br>_PMG | 0.930<br>(0.035) | **0.846**<br>(0.056) | 0.614<br>(0.027) | 0.715<br>(0.010) | **2.181**<br>(0.477) | **0.695**<br>(0.032) |
| HiMol_w/oPretrain | 0.839<br>(0.061) | 0.802<br>(0.044) | 0.580<br>(0.030) | 0.708<br>(0.014) | **3.328**<br>(1.042) | 0.815<br>(0.057) |
| HiMol_PMG | **0.898**<br>(0.028) | **0.862**<br>(0.018) | **0.602**<br>(0.030) | **0.724**<br>(0.010) | 3.459<br>(0.643) | **0.710**<br>(0.037) |

The bold values indicate the best results. $*\_PMG$ means models employ pyramid molecular graphs as model inputs instead of atomic graphs.

**Table S12.** Performance comparison evaluating the efficacy of the Pyramid Molecular Graph (PMG) as an alternative to self-supervised pre-training on the HiMol architecture.

| Models | BBBP | BACE | SIDER | ToxCast | FreeSolv | Lipophilicity |
|---|---|---|---|---|---|---|
| Himol_w/oPretrain | 0.839(0.061) | 0.802(0.044) | 0.580(0.030) | 0.708(0.014) | 3.328(1.042) | 0.815(0.057) |
| Himol_PMG | **0.898(0.028)** | **0.862(0.018)** | **0.602(0.030)** | **0.724(0.010)** | 3.459(0.643) | **0.710(0.037)** |
| Himol | 0.889(0.031) | 0.822(0.063) | 0.581(0.028) | 0.721(0.010) | **2.695(0.811)** | 0.759(0.050) |

HiMol_w/oPretrain denotes the foundational architecture without any pre-training. HiMol represents the fully pre-trained SOTA model. HiMol_PMG demonstrates our proposed variant where the native graph is replaced by the PMG topology, achieving competitive or superior results without requiring a pre-training phase.
